# Metagenomic Evidence for Horizontal Gene Transfer and Functional Convergence in the Oral Microbiome of Cohabiting Dogs and Owners

**DOI:** 10.64898/2026.04.06.716839

**Authors:** Chuantao Fang, Shan Li, Yongxiang Li, Asadullah Abid, Lu Liu, Zhuo Lan, Feng liu, Guofeng Cheng

**Affiliations:** Shanghai Tenth People’s Hospital, Institute for Infectious Diseases and Vaccine Development, Clinical Center for Brain and Spinal Cord Research of Tongji University, Affiliated Shanghai Blue Cross Brain hospital, Tongji University School of Medicine, Shanghai 200331, China; Jiangxi Provincial Key Laboratory of Cell Precision Therapy, School of Basic Medical Sciences,Jiujiang University, Jiujiang,332005, Jiangxi, China; Disease control and prevention of Canal district of Cangzhou City, Hebei, China; Digestive Endoscopy Center and department of gastroenterology, Shanghai Tenth People’s Hospital, Tongji University School of Medicine, 301 Mid. Yanchang Road, Shanghai, 200072, China

**Author notes:** Corresponding author: Guofeng Cheng,; Feng Liu. These authors contributed equally to this work (Chuantao Fang, Shan Li).

**Keywords:** Metagenome, Next Generation Sequencing, Pets, Human, Oral microbiome

## Abstract

The intimate cohabitation between humans and their pets facilitates bidirectional microbial exchange, yet the extent and functional consequences of this transfer within the oral niche remain underexplored. Here, we employed metagenomic sequencing to characterize the oral microbiome of dogs and their owners across distinct geographic regions in China, integrating taxonomic, gene-centric, and functional analyses using public databases (BacMet, CARD, eggNOG, KEGG) to assess microbe–host associations. We found that dog–owner pairs exhibited significantly higher gene-level similarity compared to unrelated individuals, indicating a strong cohabitation-driven microbial linkage. While no major taxonomic shifts were observed in the human oral microbiome associated with pet ownership, we identified a marked enrichment of antibiotic resistance genes (ARGs)—particularly those conferring resistance to peptides, fluoroquinolones, antiseptics, diaminopyrimidines, cephalosporins, and carbapenems—in cohabiting pairs. This enrichment, together with the identification of exclusively shared ARGs (e.g., mdtF, macB, RanA), suggests the potential for horizontal gene transfer (HGT) between pet- and human-associated microbiomes. Functional profiling further revealed greater similarity in microbial metabolic pathways between cohabiting pairs than between unrelated individuals, reinforcing the likelihood of HGT as a mechanism underlying functional convergence. Collectively, these findings reveal that cohabitation with dogs reshapes the human oral microbiome at the genetic and functional levels, with potential implications for antimicrobial resistance transmission. This study provides a foundational framework for assessing the health risks associated with pet–human microbial exchange in shared living environments.

## Introduction

The microbiome comprises trillions of microorganisms that live in a symbiotic fashion with their host [1]. They can exist as commensals or pathogens, and their composition depends on the host’s diet, lifestyle, and external environment[2]. It represents a complex, dynamic system comprising bacteria, fungi, viruses, archaea, and microbial eukaryotes, which is critical to human and animal life[3]. The gut microbiome plays a pivotal role in host physiology, contributing to the biosynthesis of essential vitamins, modulation of immune responses against pathogenic microorganisms, and maintenance of nutritional homeostasis and overall host health [3, 4]. Additionally, it confers colonization resistance, thereby inhibiting the proliferation of invading pathogens [5]. Perturbations in the composition and function of the gut microbiota, a state termed dysbiosis, can precipitate the overgrowth of specific microbial taxa [6, 7] and predispose the host to various disease etiologies [8]. Such dysbiotic states have been implicated in critical public health concerns, including the transmission of epidemic diseases via companion animals. Consequently, a resilient and well-balanced gut microbiome is paramount for preserving host health and mediating protection against opportunistic and pathogenic microorganisms.

The oral cavity, serving as a primary gateway for food ingestion, harbors a highly dense and diverse microbiome. It encompasses multiple microbial niches (including the teeth, tongue, hard and soft palate, gingiva, and cheeks), which collectively constitute a unique ecosystem [9]. The normal oral microflora consists of a multitude of microorganisms that exhibit either protective or pathogenic properties [10]. The composition of the oral microflora is modulated by a multitude of factors, including age, dietary intake, environmental fluctuations, and host health status. When host health deteriorates, certain oral microorganisms disseminate to internal organs, leading to opportunistic infections [10]. During dysbiosis, this ecosystem creates a permissive environment for pathogenic microorganisms, contributing to a spectrum of acute and chronic diseases, including periodontal disease [9, 11]. Therefore, it is meaningful to capture the microorganism profiles of the oral cavity.

Pets are common companion animals that maintain intimate contact with their owners, consequently exchanging microbial communities with humans [10]. A subset of these microbes is zoonotic, capable of inducing disease in humans, including bacterial pathogens (e.g., *Campylobacter*, *Salmonella*), viral agents (e.g., rabies), dermatophytic fungi, and protozoan parasites (e.g., *Giardia*, *Cryptosporidium*) [12, 13]. These pathogens elicit clinical manifestations ranging from mild cutaneous rashes to severe neurological symptoms in humans [12]. Furthermore, Pets can acquire pathogens such as influenza viruses, beta-lactamase-resistant *Escherichia coli*, and *Clostridium difficile*, which pose significant risks to their owner’s health [12]. Researchers are actively investigating shifts in microbial species associated with host health or disease states [4]. Elucidating these dynamics is critical for deciphering the pet-human relationship, particularly with respect to how their interactions modulate microbial composition and impact health outcomes.

In this study, we employed metagenomic approaches to characterize the diversity of the oral microbiome in dogs and their owners across different regions of China. We aimed to develop an integrated and accessible framework that highlights shared microbiome profiles, thereby opening future research avenues and facilitating the establishment of more efficient healthcare systems.

## Methods

### Sample collection

Oral swabs were collected from volunteers (N = 6×10), dog owners (N = 8×10), and dogs (N = 8×10) via convenience sampling under sterile conditions. Samples were obtained from eight eastern provinces of China (including Shanghai, Dalian, Tianjin, Shandong, Guangdong, Jiangxi, Hebei, and Guizhou), with 30 samples collected from each province. Only participants (humans and dog owners) and dogs that had abstained from antibiotic use for at least 60 days before sample collection were included in the study. The study was approved by the ethical committee of the School of Medicine of Tongji University (approved #: 2023tjdxsy031). Oral swab collection followed the method described by Lisjak et al [14]. Briefly, a single swab was rotated throughout the entire oral cavity (including the tongue, cheeks, and buccal mucosa) to capture the comprehensive oral microbiome of each individual. These samples were immediately placed on ice, transported to the School of Medicine, Tongji University, and stored at -80 °C until subsequent nucleic acid extraction.

### Librarying

We utilized a pooling strategy wherein 10 samples per region were combined into a mixed sample. For metagenomic analyses, total genomic DNA was isolated using the ZymoBIOMICS DNA Kit (Zymo Research, Irvine, CA, USA) in strict accordance with the manufacturer’s protocols. Before library construction, sample concentration, integrity, and purity were assessed using the Qubit Fluorometer (Thermo Fisher Scientific, Waltham, MA, USA) and NanoDrop Spectrophotometer (Thermo Fisher Scientific, Waltham, MA, USA), respectively. Metagenomic library construction and sequencing were performed as described previously [15] with minor modifications. Briefly, 500 ng of genomic DNA was sheared into ∼350 base pairs (bp) fragments using a Covaris E220 Focused-ultrasonicator (Covaris, Brighton, UK) at 175 W peak incident power, 10% duty factor, 200 cycles per burst, and 80 seconds of treatment. End repair and A-tailing were carried out sequentially: the sheared fragments were repaired to generate blunt ends, followed by the addition of adenine (A) residues to the 3’ ends to facilitate adapter ligation. Adapters were ligated to the A-tailed fragments using the MGIEasy DNA Adapter Kit (MGI Tech, Shenzhen, China) per the manufacturer’s instructions.

Polymerase chain reaction (PCR) was employed to amplify the adapter-ligated fragments using a high-fidelity Kapa HiFi DNA polymerase (Roche, Basel, Switzerland) with 12 cycles of amplification (98°C for 20 seconds, 60°C for 30 seconds, 72°C for 30 seconds). Amplification products were purified using AMPure XP magnetic beads (Beckman Coulter, Brea, CA, USA) at a 1.8× ratio to remove short fragments (<150 bp) and residual primers. The purified PCR products were denatured at 95°C for 3 minutes and circularized using a thermostable ligase to generate single-stranded circular DNA libraries, a key step for optimizing DNBSEQ sequencing efficiency.

### Sequencing

Libraries were normalized to a concentration of 10 nM using the Qubit Fluorometer, pooled in equal volumes, and converted into DNA nanoballs (DNBs) via rolling-circle amplification using the DNBSEQ Library Prep Kit (MGI Tech, Shenzhen, China). The pooled DNBs were loaded onto a DNBSEQ-G400 flow cell and sequenced in paired-end 150 bp mode (PE150) on a DNBSEQ-G400 sequencer (MGI Tech, Shenzhen, China) in strict adherence to the manufacturer’s specifications for read length, cluster density, and sequencing chemistry.

### Gene Assembly

After the acquisition of raw sequence reads, whole-genome analysis (WGA) was executed utilizing BGI’s proprietary bioinformatics pipeline. Briefly, high-quality clean reads were generated via SOAPnuke software (https://github.com/BGI-flexlab/SOAPnuke) through the removal of low-quality reads, adapter sequences, and ambiguous bases from the dataset. These reads were aligned with the Bowtie2 algorithm (https://bowtie-bio.sourceforge.net/bowtie2/index.shtml) to exclude host-derived reads, employing the host genome as a reference (Human, https://ftp.ensembl.org/pub/release-115/fasta/homo_sapiens/dna/Homo_sapiens.GRCh38.dna.toplevel.fa.gz; Dog, https://ftp.ensembl.org/pub/release-115/fasta/canis_lupus_familiaris/dna/Canis_lupus_familiaris.ROS_Cfam_1.0.dna.toplevel.fa.gz). The filtered reads were assembled into contigs using MEGAHIT (https://github.com/voutcn/megahit), followed by the exclusion of fragments shorter than 300 bp. Contig continuity and integrity were evaluated by computing metrics including total assembly length, minimum and maximum contig lengths, and N50/N90 values.

### Gene expression quantification

Gene sequences within the assembled contigs were predicted using MetaGeneMark (https://github.com/gatech-genemark/MetaGeneMark-2), followed by the deduplication of the resulting gene list via CD-HIT (https://www.bioinformatics.org/cd-hit/) using a predefined sequence similarity threshold. Relative gene abundances were quantified using Salmon, a tool optimized for transcript-level quantification. Protein-coding sequences (CDSs) were predicted via an open reading frame (ORF) finder to construct a high-quality gene catalog. Alpha- and beta-diversity analyses of the genetic data were conducted using the Vegan R package to assess compositional variation.

### Functional annotation

Functional annotation of non-redundant genes was performed using eggNOG-mapper (originally referenced as DIAMOND; https://github.com/bbuchfink/diamond) against multiple databases, including the Kyoto Encyclopedia of Genes and Genomes (KEGG), Bacterial Antibacterial Biocide & Metal Resistance Genes Database (BacMet), Comprehensive Antibiotic Resistance Database (CARD), etc. KEGG was stratified into three hierarchical levels for downstream analysis.

Taxonomic classification from higher to lower hierarchical levels was assigned using Kraken2 (for initial classification) and Bracken2 (for abundance estimation). Taxonomic alpha- and beta-diversity indices were calculated based on species-level relative abundances. Alpha diversity quantifies within-sample or within-community diversity, encompassing both species richness (number of distinct species, Chao1 index) and evenness (relative abundance of species, Shannon and Simpson index). Beta diversity, by contrast, assesses between-sample or between-community differences in species composition, in which three independent methods (including Jensen-Shannon, Euclidean, and Bray-Curtis) were used to estimate the distance.

### Statistical analysis

Alpha diversity indices were evaluated via the nonparametric Kruskal-Wallis test implemented in GraphPad Prism v10.3. When overall results reached statistical significance, post-hoc pairwise comparisons were performed using Dunn’s multiple comparison test. Statistical significance was defined as p < 0.05. In the case of multiple testing (for beta diversity), ADONIS (Permutational Multivariate Analysis of Variance Using Distance Matrices) and ANOSIM (Analysis of Similarities) were used, and adjusted p-values (P-adj) were computed using the Benjamini-Krieger-Yekutieli method to control the false discovery rate (FDR).

## Results

### Microbiomes revealed that pet and pet owner groups exhibit greater gene-level similarity

To investigate different microbiomes between pets and their owners, we conducted metagenomic sequencing on oral swabs from dogs (n = 8), their owners (n = 8), and unrelated volunteers (n = 6), which were collected from 8 cities/provinces of China, covering almost 240 individuals (**Fig. 1A and B**). After quality control (including adapter sequence trimming, removal of low-quality reads, and elimination of host genomic DNA contaminants), we obtained over 1.6 billion high-quality reads from 22 libraries, exhibiting balanced GC contents, with no significant inter-group variation, indicative of robust sequencing quality and reliable downstream analytical potential (**Supplementary Tables S1–S3**).

**Fig. 1.**
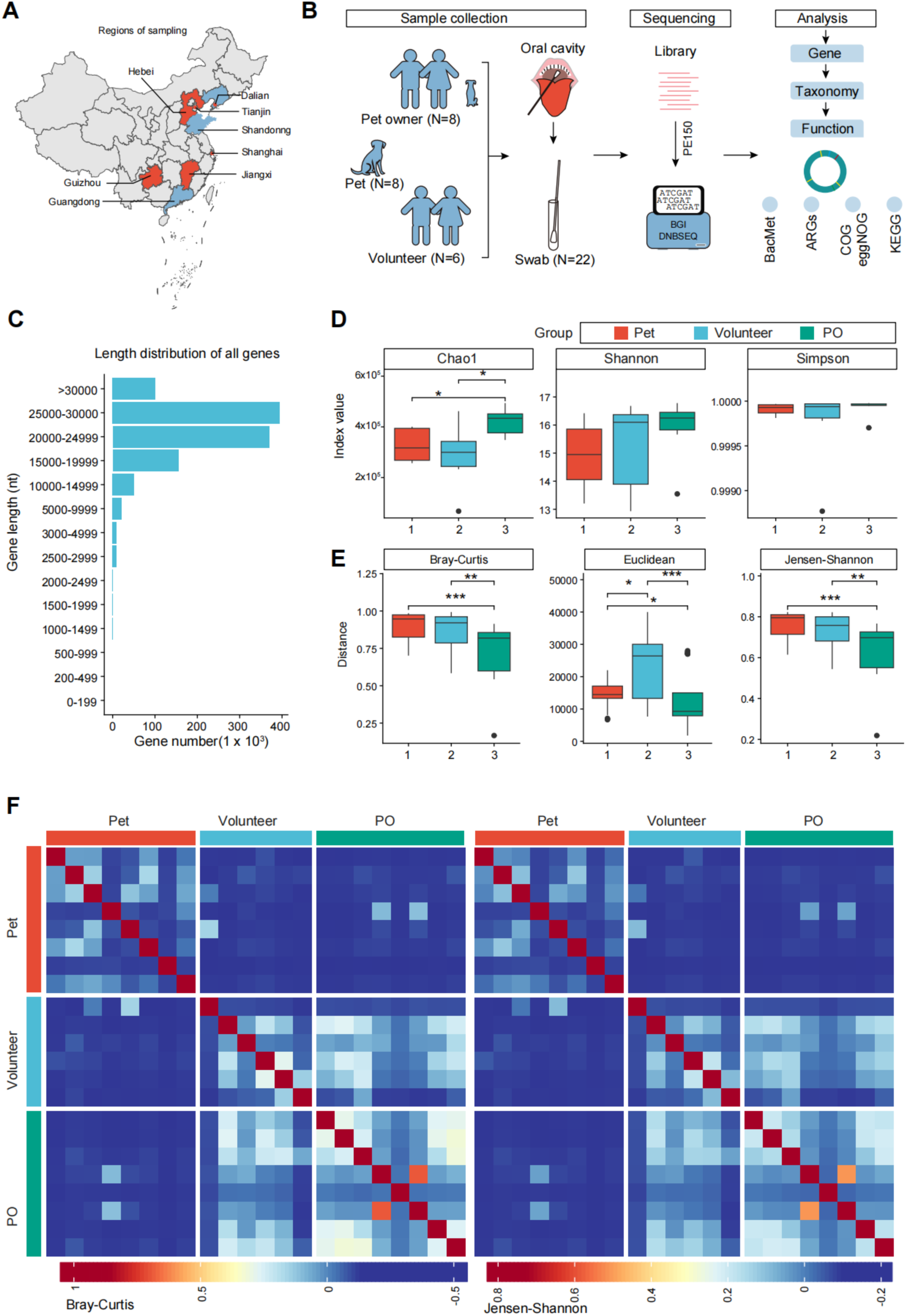
Microbiome analysis of dogs and their owners at gene-level. **A.** Geographical distribution of the sampling sites from various regions of China. Analytic flow of the current study. PO, pet owner. BacMet, Antibacterial Biocide & Metal Resistance Genes Database; ARGs, Antibiotic resistance genes; COG, Clusters of Orthologous Genes database; eggNOG, a public database of orthology relationships, gene evolutionary histories, and functional annotations; KEGG, Kyoto Encyclopedia of Genes and Genomes. **B.** Gene length distribution across all samples. **C.** Chao1/Shannon/Simpson index representing the overall richness and evenness of microbial species across experimental groups. **D.** Distance of microbial community’s functional gene composition. Distances were calculated by Bray-Curtis, Euclidean, and Jensen-Shannon methods, respectively. Each point represents a sample, colored according to its group. Group volunteer (N = 6), dog owner (N = 8), and dog (N = 8). **E.** The stars show the level of significance *p < 0.05, **p < 0.01, ***p < 0.001. **F.** Heatmap showing the clustering of the indicated samples from different groups. Distances were calculated by Bray-Curtis, Jensen-Shannon method, respectively.

Following de novo assembly, the resulting genes possessed lengths exceeding 1000 bp, with the majority distributed between 20–30 kb (**Fig. 1C and Table S2**), reflecting efficient assembly of diverse microbial genomic fragments. The different samples exhibited a huge variation in the assembly contig numbers (from ∼2000 to 200,000) (**Supplementary Tables S2**), showing an obvious heterogeneity among samples. To systematically characterize the global gene-level features of all samples, we conducted alpha-diversity analyses using three ecological indices: Chao1 (for microbial richness), Shannon (for combined diversity and evenness), and Simpson (for dominance) on all recovered genes. We observed that the total number of recovered genes varied significantly among groups (**Fig. 1D and Supplementary Tables S3**), with dog and dog owner groups harboring a greater number of genes than volunteer groups (**Fig. 1D**, Chao1 index). However, recovered genes from different groups displayed comparable levels of abundance, evenness, and diversity (**Fig. 1D**, Shannon and Simpson indices). These results indicate that while cohabiting dogs and their owners exhibit greater microbial richness, the structural complexity and uniformity of their oral microbiomes are analogous among the three groups.

To further investigate inter-group structural similarities and dissimilarities, we performed beta-diversity analyses using Bray-Curtis dissimilarity indices, which quantify taxonomic and functional distances between microbial communities. Interestingly, genes from dog and dog owner groups exhibited a degree of similarity and were significantly distinct from those of volunteer groups (**Fig. 1E**). Consistently, dog samples clustered more closely with dog owner groups than with volunteer groups (**Fig. 1F**). Altogether, our findings demonstrate that dog and dog owner groups exhibit greater gene-level similarity than volunteer groups, indicating a closer microbial association between pets and their owners.

### Significant alterations at taxonomic levels associated with dog ownership

We further performed taxonomic analysis on the obtained sequencing data, and alpha- and beta-diversity analysis (Shannon, Simpson, and Chao1 for alpha; Bray-Curtis for beta-diversity), taxonomic components were carried out to estimate the difference among the three groups. Our analysis included six taxonomic ranks: phylum, class, order, family, genus, and species levels. For alpha-diversity metrics, the Chao1 index revealed significant differences in total species count exclusively between the dog and volunteer cohorts at the order, family, genus, and species levels (**Fig. 2A and Supplementary Fig. S1A**). In contrast, assessments using the Shannon and Simpson indices, which integrate both species richness and evenness, demonstrated that variations in species diversity and richness were primarily evident between the dog and dog owner groups at the phylum and class levels (**Fig. 2A and Supplementary Fig. S1A**). Notably, beta-diversity analyses based on Bray-Curtis dissimilarity revealed significant intergroup distinctions in both species composition and relative abundance across all three cohorts, with the most pronounced differences detected at the species level (**Fig. 2B and Supplementary Fig. S1B**). Complementary principal coordinate analysis (PCoA) further corroborated this partitioning, showing that dog-derived samples formed distinct clusters and were phylogenetically distant from samples collected from the other two groups across all six taxonomic ranks (**Supplementary Fig. S1B**).

**Fig. 1.**
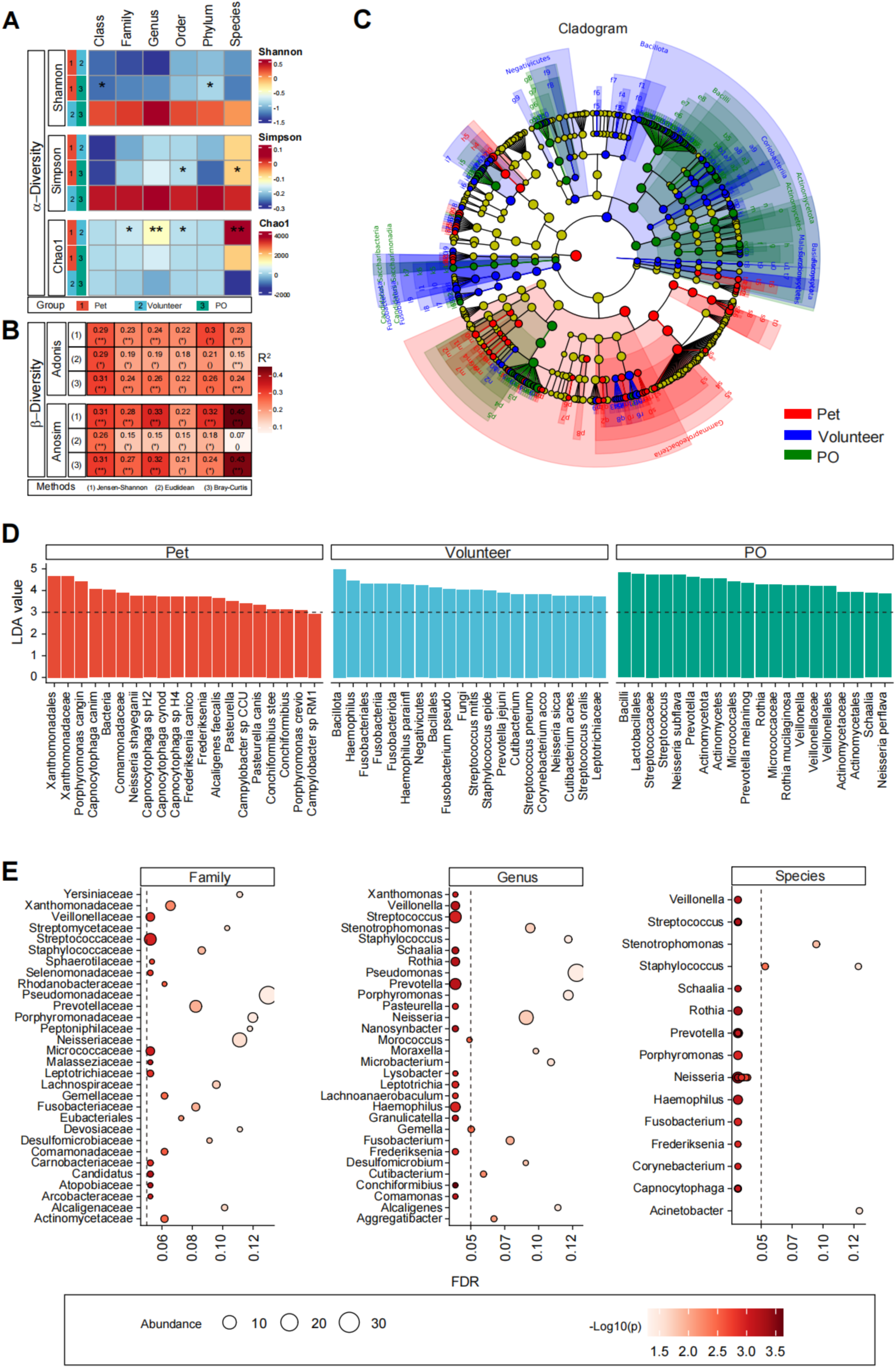
Significant alterations at taxonomic levels associated with dog ownership. **A.** Heatmap showing the comparison results of alpha-diversity of taxonomic components among Volunteer, PO (pet owner), and dog groups. Taxonomic annotations were carried out at the Phylum, Class, Order, Family, Genus, and Species levels. The taxonomic distance of each sample was reflected by the Chao1/Shannon/Simpson index. *p < 0.05, **p < 0.01, ***p < 0.001. **B.** Heatmap showing the comparison results of beta-diversity of taxonomic components among Volunteer, DO, and dog groups. The taxonomic distance among samples in each group was reflected by the Bray-Curtis and Jensen-Shannon methods, respectively. Adonis and Anosim methods were used for statistical analysis. *p < 0.05, **p < 0.01, ***p < 0.001. **C.** LEfSe analysis revealing the taxa with significant differences associated with dog ownership. **D.** Barplot showing the differentially expressed taxonomic components. The dashed line shows the cutoff (LDA > 3). LDA, the value of Linear Discriminant Analysis. **E.** Dotplot showing the differentially expressed taxonomic components among Volunteer, PO (pet owner), and dog groups. The dashed line shows the cutoff (p <0.05).

While overall changes in community structure were significant, our analysis revealed significant alterations at lower taxonomic levels associated with dog ownership (**Supplementary Fig. S2A**). Notably, substantial inter-individual variability was observed in the magnitude of differential species abundance across the study cohort (**Supplementary Fig. S2A**). Linear discriminant analysis effect size (LEfSe) was employed to identify taxonomic biomarkers distinguishing the study groups: individuals not cohabiting with dogs were enriched in commensal taxa characteristic of a healthy human oral cavity, including *Bacillota*, *Prevotellaceae*, *Haemophilus*, *Fusobacteriales*, *Fusobacteriia*, and *Haemophilus parainfluenzae* (LDA > 4) (**Fig. 2C and D**). In contrast, the dog owner group was characterized by a predominance of *Neisseria*, *Neisseriaceae*, *Neisseriales*, *Bacilli*, *Lactobacillales*, and *Streptococcus* (LDA > 4) (**Fig. 2C and D**). Many of these enriched taxa in dog owners are well-characterized components of human-adapted mucosal microbial communities, thereby indicating a pattern of microbiota convergence between cohabiting humans and their canine companions—likely driven by sustained close contact in shared domestic environments (**Fig. 2C and D**). The taxonomic biomarkers reveal a clear partitioning of microbial lineages across experimental groups, reflecting both host specificity and cross-exposure signatures. Additionally, several genera were differentially distributed among the three groups (**Fig. 2E**). Altogether, no consistent directional shifts in the overall human oral microbiome were observed at higher taxonomic ranks in cohabiting domestic environments, despite the fine-scale taxonomic alterations linked to dog ownership.

### Antibiotic resistance genes reveal potential horizontal gene transfer between dogs and their owners

Antibiotic resistance genes (ARGs) are genetic elements encoding specific defense mechanisms that confer upon microorganisms the ability to survive exposure to antimicrobial agents. As a component of microbial evolutionary adaptation, ARGs are naturally ubiquitous across a wide range of terrestrial, aquatic, and host-associated ecosystems, representing ancient evolutionary traits shaped by long-term microbial-antimicrobial interactions [16]. Critically, ARGs can spread via horizontal gene transfer (HGT) through mobile genetic elements (including integrons and plasmids), facilitating cross-species transmission between pathogenic and non-pathogenic bacteria, such as between pets and their owners. Given the World Health Organization’s (WHO) warning of rising mortality from resistant infections, ARGs are recognized as a foundational and primary driver of the ongoing global antibiotic resistance crisis [17].

In the present study, we conducted systematic functional annotation and comparative analyses of acquired protein-coding sequences associated with antimicrobial resistance. In total, we identified 1238 ARGs, which were categorized into 27 distinct classes, including those conferring resistance to minocoumarin, aminoglycosides, fluoroquinolones, tetracyclines, and multiple other major antimicrobial categories (**Fig. 3A and Supplementary Fig. S3A**). Notably, while the total abundance of ARGs was broadly uniformly distributed across all experimental groups, marked inter-individual variability was observed within each group (**Supplementary Fig. S4A**). Concurrently, hierarchical clustering and relative abundance profiling revealed striking similarities in both the overall compositional structure and quantitative abundance of ARGs between the dog and dog owner cohorts **(Fig. 3A and B**). However, total ARG abundance (copies/cell) exhibited no difference among the three groups (**Fig. 3C**). Intriguingly, ARGs belonging to the peptide, fluoroquinolone, antiseptic, diaminopyrimidine, cephalosporin, and carbapenem resistance classes displayed significantly higher richness in both the dog and dog owner cohorts relative to the control group (**Fig. 3D and Supplementary Fig. S4B**), suggesting potential HGT between pets and their owners. Consistently, we further identified a subset of ARGs (including mdtF, macB, and RanA) that were exclusively detected in paired samples from companion animals and their corresponding owners (**Fig. 3E**).

**Fig. 3.**
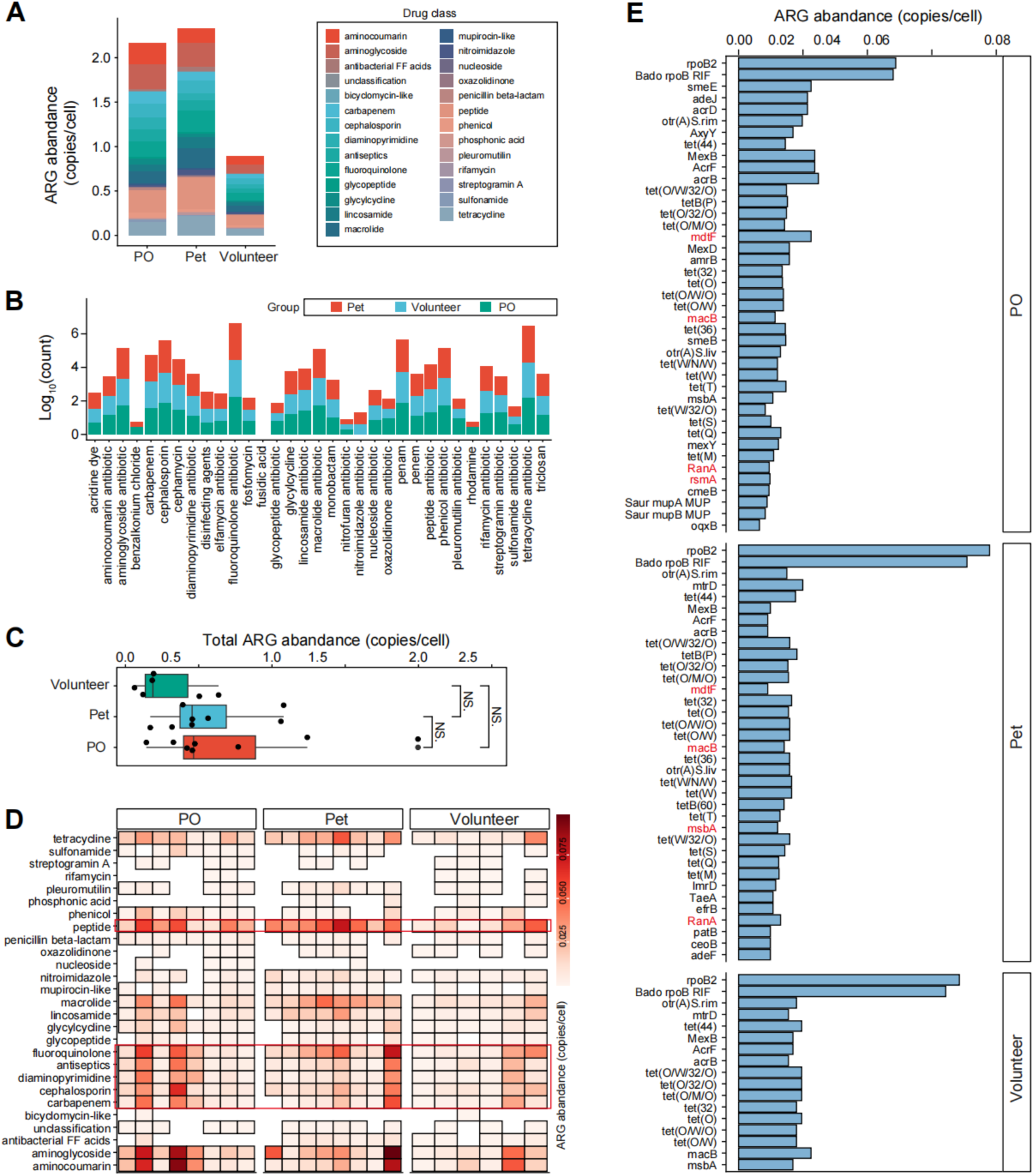
Horizontal gene transfer (HGT) of antibiotic resistance genes between dogs and their owners. **A.** Boxplot showing the relative abundance of ARG components of Volunteer, PO (pet owner), and dog groups. PO, pet owner; ARGs, Antibiotic resistance genes. **B.** Barplot showing the total counts of ARG components of Volunteer, PO (pet owner), and dog groups. PO, pet owner; ARGs, Antibiotic resistance genes. **C.** Boxplot showing the total abundances of ARG components (per cell) of Volunteer, PO (pet owner), and dog groups. PO, pet owner; ARGs, Antibiotic resistance genes. NS, no significance in statistics. **D.** Heatmap showing the details of ARG components (per cell) of Volunteer, PO (pet owner), and dog groups. PO, pet owner. **E.** Barplot showing the differentially expressed ARG components.

### BacMet annotation exhibited most pronounced differences observed between pets and dog owners

Antibacterial biocides and heavy metals are well-documented to drive the selection of antibiotic-resistant bacterial populations via characterized co-resistance and cross-resistance mechanisms—whereby exposure to one selective agent can confer unintended resistance to structurally or functionally unrelated antimicrobials or toxic compounds. Notably, these resistance traits can be horizontally transferred, enabling their co-transmission between companion animals and their human owners in shared household microenvironments. In the present study, we leveraged the curated BacMet database to systematically identify, functionally annotate, and quantify the prevalence and relative abundance of biocide resistance genes (BMRGs) and metal resistance genes (MRGs) within host-associated microbial communities [18, 19], thereby uncovering links between specific microbes and environmental contaminants.

Overall, BMRGs/MRGs exhibited significant variation across dogs, dog owners, and volunteer participants, with the most pronounced differences observed between pets and dog owners (**Supplementary Fig. S3A and B**). Collectively, a total of 31 unique BMRG/MRG subtypes were identified across all cohorts, with the relative proportional distribution of BacMet functional categories demonstrating only minimal heterogeneity across pet dogs, volunteers, and owners (**Fig. 4A**). Notably, among the identified BMRGs/MRGs, the most highly abundant subtypes encoded resistance mechanisms targeting peroxides, phenolic compounds, acridine derivatives, and several other structurally and functionally distinct antimicrobial chemical classes (**Fig. 4B**). Subsequent differential abundance analysis of BMRGs/MRGs across the three cohorts revealed statistically significant, group-specific enrichment of genes mediating resistance to antimony, arsenic, chlorhexidine, ethidium bromide, and a panel of additional antimicrobial and heavy metal compounds (**Fig. 4C**). More importantly, comparisons between pets and dog owners accounted for the majority of the variability, with prominent differences observed in genes linked to arsenic, ethidium bromide, rhodamine 6G, tellurium, and other agents (**Fig. 4D**). Notably, pets and dog owners exhibited a higher degree of shared BMRGs/MRGs compared to other groups (**Fig. 4E**). Overall, the results integrate compositional, quantitative, and correlative analyses to delineate BacMet profiles in pets and humans, thereby emphasizing group-specific patterns and gene-host relationships.

**Fig. 4.**
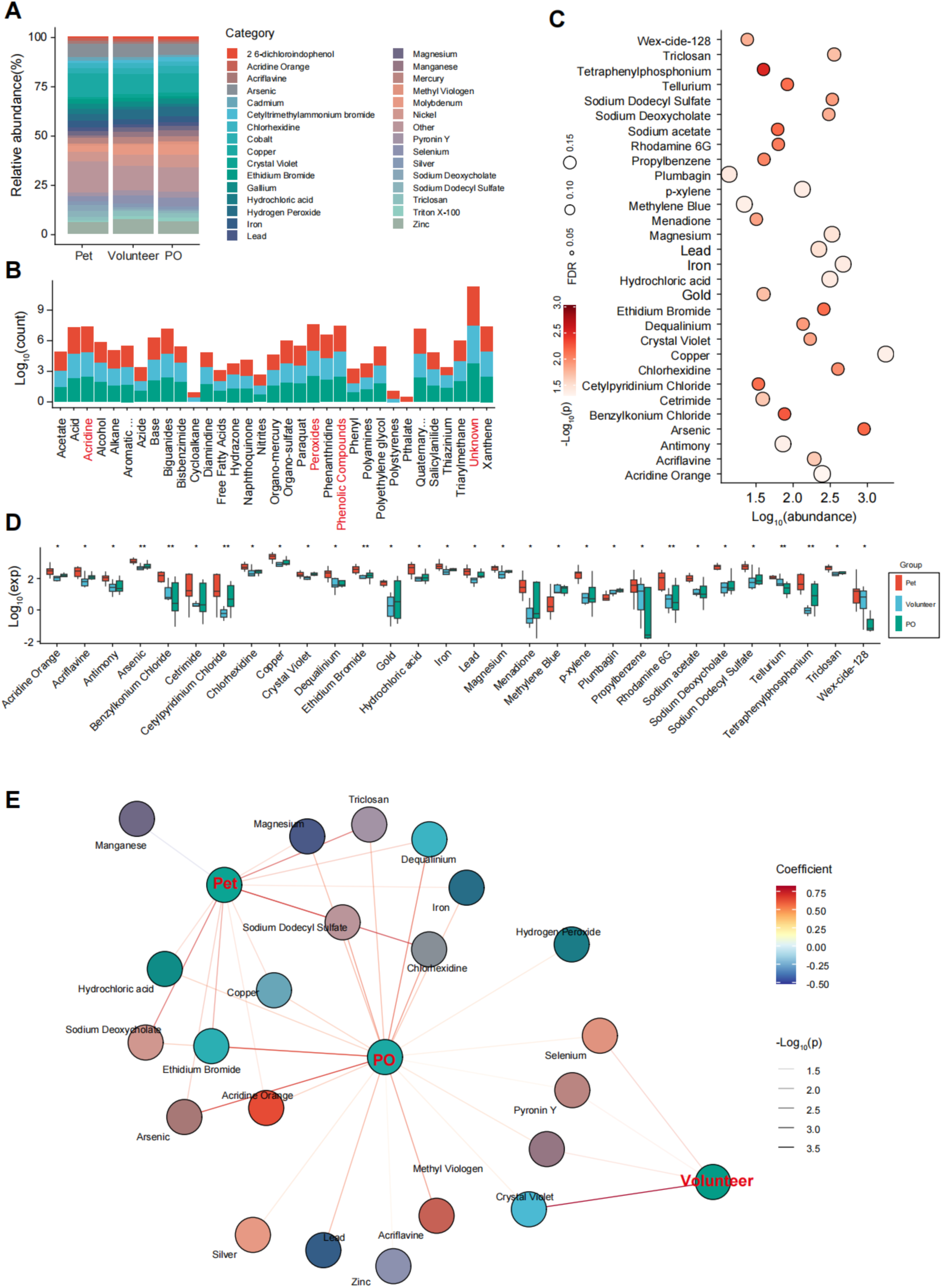
BacMet annotation analysis between dogs and their owners. **A.** Boxplot showing the relative abundance of BacMet components of Volunteer, PO (pet owner), and dog groups. PO, pet owner; BacMet, Antibacterial Biocide & Metal Resistance Genes Database. **B.** Barplot showing the total counts of BacMet components of Volunteer, PO (pet owner), and dog groups. **C.** Dotplot showing the differentially expressed BacMet components among Volunteer, PO (pet owner), and dog groups. **D.** Boxplot showing the total abundances of BacMet components of Volunteer, PO (pet owner), and dog groups. *p < 0.05, **p < 0.01, ***p < 0.001 by Kruskal-Wallis test. **E.** The functional correlation analysis of BacMet components.

### COG/eggNOG annotation revealed HGT events occurring between domestic pets and their human cohabitants

To gain deeper insights into the functional roles of genes and proteins across divergent species [20], we conducted systematic functional annotation of the sequencing data generated in this study, leveraging curated reference datasets from well-established orthologous repositories, including the Clusters of Orthologous Genes (COG) database and the evolutionary genealogy of genes: Non-supervised Orthologous Groups (eggNOG) database. This annotation facilitates the interpretation of genomic data and supports various applications in microbial genomics (including microbial genome annotation and comparative genomics) through the classification of genes and proteins into evolutionarily defined orthologous groups [21–23].

In the COG annotation analysis, orthologous gene groups displayed substantial inter-group variation among canine subjects, their cohabiting owners, and unrelated volunteer participants. Notably, the most striking disparities were detected between canine pets and human participants (both owners and volunteers) (**Supplementary Fig. S3A and B**). A total of 24 distinct functional categories were assigned to the retrieved coding sequences via COG annotation, including amino acid transport and metabolism, carbohydrate transport and metabolism, cytoskeleton, cell motility, and cell wall/membrane/envelope biogenesis, which displayed nearly balanced abundances across the three groups (**Fig. 5A and B**). Similarly, 23 functional categories were detected in eggNOG annotation, encompassing amino acid transport and metabolism, carbohydrate transport and metabolism, cell motility, cytoskeleton, and translation/ribosomal structure and biogenesis (**Supplementary Fig. S5A and B**). Subsequent differential functional analysis revealed that several key functional categories—specifically cell wall/membrane/envelope biogenesis, energy production and conversion, and signal transduction mechanisms—showed more pronounced inter-group divergence across the three cohorts (**Fig. 5C and S5C**). Most critically, our comparative analysis demonstrated that canine subjects shared a significantly higher number of homologous functional categories with their respective owners than with unrelated volunteer participants, strongly suggesting HGT events occurring between domestic pets and their human cohabitants (**Fig. 5D and Supplementary Fig. S5D**).

**Fig. 5.**
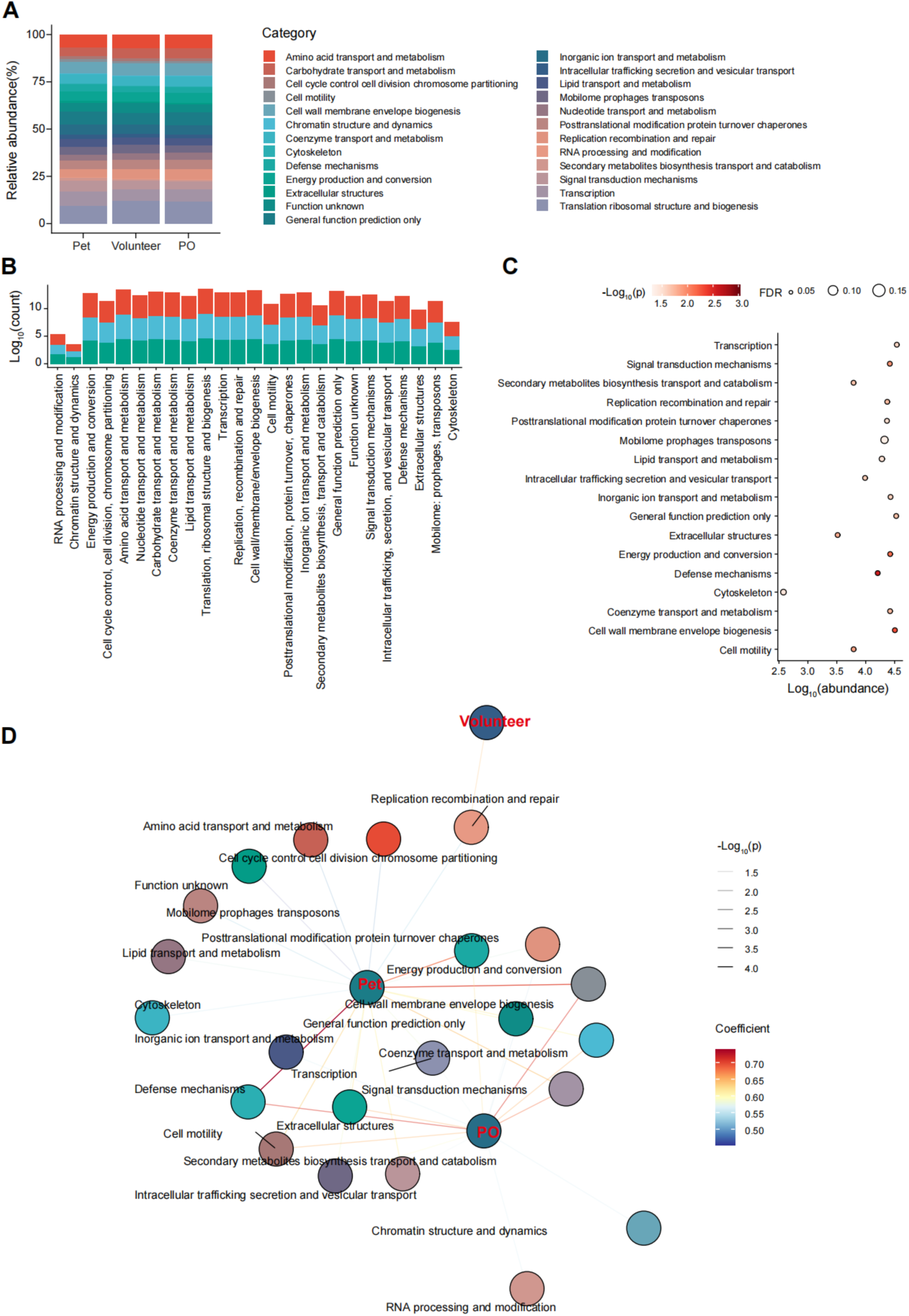
COG/eggNOG annotation revealed HGT events occurring between dogs and their owners. **A.** Boxplot showing the relative abundance of COG components of Volunteer, PO (pet owner), and dog groups. PO, pet owner; COG, Clusters of Orthologous Genes database. **B.** Barplot showing the total counts of COGt components of Volunteer, PO (pet owner), and dog groups. **C.** Dotplot showing the differentially expressed COG components among Volunteer, PO (pet owner), and dog groups. **D.** The functional correlation analysis of COG components.

### KEGG analysis supported greater functional similarity between the dog and dog owners

Besides, we performed Kyoto Encyclopedia of Genes and Genomes (KEGG) annotation on the obtained coding sequences, which assigns functional annotations to genes, proteins, and other biological molecules by leveraging resources from the KEGG database, thereby facilitating systematic analysis of gene functions and metabolic/signaling pathways in biological systems [24]. Overall, global comparative analysis of KEGG pathway profiles revealed a significantly higher degree of divergence in the dog group relative to the other two experimental groups (**Supplementary Fig. S3A and B**). Collectively, a total of 28 Level 2 and 31 Level 3 KEGG pathways were identified across the translated protein sequences of all three groups, including 1) amino acid metabolism, 2) biosynthesis of other secondary metabolites, 3) antimicrobial drug resistance, 4) environmental adaptation, 5) glycan biosynthesis and metabolism, 6) bacterial infectious disease, 7) metabolism of cofactors and vitamins, etc (**Fig. 6A and Supplementary Fig. S6A-B**). Concurrently, we observed that these pathways were almost uniformly distributed across the groups (**Fig. 6A and Supplementary Fig. S6A**). Most prominently, metabolic pathways constituted the most abundant category in all groups, representing the largest proportion of functionally annotated pathways across each experimental cohort (**Supplementary Fig. S6A**).

**Fig. 6.**
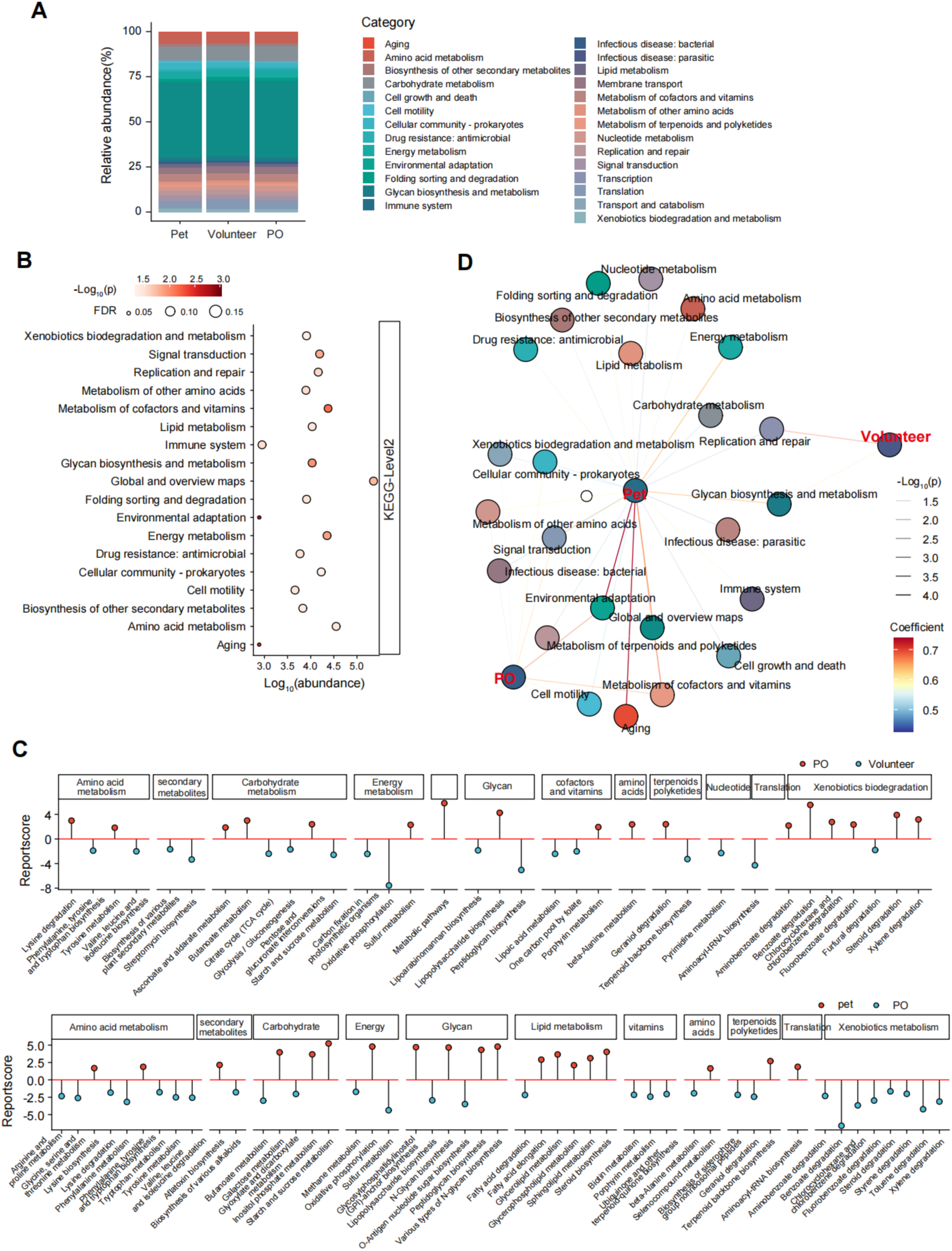
KEGG analysis supported greater functional similarity between the dog and dog owners. **A.** Boxplot showing the relative abundance of KEGG components of Volunteer, PO (pet owner), and dog groups. PO, pet owner; KEGG, Kyoto Encyclopedia of Genes and Genomes. **B.** Dot plot showing the differentially expressed KEGG components among Volunteer, PO (pet owner), and dog groups. **C.** Lollipop chart showing the differentially expressed KEGG components of the indicated comparisons. **D.** The functional correlation analysis at KEGG components.

In the differential analysis, we identified that pathways associated with cofactor and vitamin metabolism, glycan biosynthesis and metabolism, as well as energy metabolism displayed pronounced inter-group variability across the three cohorts (**Fig. 6B**). At the KEGG level 3 functional annotation, specifically, pathways including carbon metabolism, O-antigen nucleotide sugar biosynthesis, the citrate cycle (TCA cycle), and cofactor biosynthesis were found to be significantly differentially represented across groups (**Supplementary Fig. S6C**). In the comparison of the volunteer and dog owner groups, a greater number of xenobiotic biodegradation terms (including aminobenzoate degradation, benzoate degradation, fluorobenzoate degradation, steroid degradation, and xylene degradation) were enriched in the dog owner group (**Fig. 6C**). Conversely, the volunteer cohort exhibited significant enrichment of pathways involved in secondary metabolite biosynthesis, energy metabolism, nucleotide metabolism, and translational processes (**Fig. 6C**). In the comparison of the dog and dog owner groups, the dog group was enriched for lipid metabolism and glycan-related pathways, including fatty acid elongation, glycerolipid metabolism, glycerophospholipid metabolism, sphingolipid metabolism, steroid biosynthesis, N-glycan biosynthesis, peptidoglycan biosynthesis, various N-glycan biosynthesis pathways, etc (**Fig. 6C**). Notably, we observed greater similarity between the dog and dog owner groups than between either group and the volunteer group (**Fig. 6D and Supplementary Fig. S6D**).

## Discussion

The interaction between humans and pets impacts microbial ecosystems in shared environments. Understanding the dynamics of microbial populations is crucial to elucidating health outcomes, particularly with respect to how microbial composition influences such outcomes. We investigated the effects of cohabitation on microbial population dynamics using a metagenomic approach, elucidating the diversity, composition, and functional activity of microbial communities across human and their pets. Our study revealed distinct microbial functional profiles among dogs, their owners, and volunteers, with greater functional overlap between pets and owners than with volunteers, as evidenced by BacMet, CARD, COG, eggNOG, KEGG annotations, etc. These findings provide novel insights into the functional interactions of the human-dog microbiome and potential horizontal gene transfer driven by cohabitation.

In the analysis of metagenomic profiles, we observed that the variation in orthologous group classification across groups (particularly the pronounced differences between pets and humans) aligns with previous comparative genomics studies [25]. The greater functional similarity between pets and owners compared to volunteers supports the hypothesis of HGT, a phenomenon previously implicated in human-pet microbiome interactions [26, 27]. For instance, the enrichment of cell wall/membrane biogenesis and signal transduction in differential analysis may reflect adaptive responses to shared environmental pressures (e.g., diet, household chemicals), where HGT could enhance microbial fitness in both hosts. However, the indirect nature of our evidence (relying on functional overlap rather than direct gene transfer validation) highlights the need for follow-up studies using metagenome-assembled genomes to identify specific transferred gene fragments.

The metabolic pathways, such as amino acid metabolism, biosynthesis of other secondary metabolites, antimicrobial drug resistance, environmental adaptation, etc were dominant in all groups, being consistent with the primary role of gut microbiota in nutrient processing [28, 29]. The enrichment of xenobiotic biodegradation pathways in owners (e.g., aminobenzoate, benzoate degradation) likely stems from their higher exposure to environmental pollutants (e.g., personal care products, pesticides), a finding supported by studies linking human lifestyle to enhanced microbial xenobiotic metabolism [30, 31]. In contrast, the enrichment of lipid and glycan biosynthesis pathways in pets may reflect their dietary habits (e.g., high-fat commercial diets) or inherent microbial adaptations to canine oral cavity physiology, as reported in previous canine microbiome studies [32, 33]. The greater similarity in KEGG profiles between pets and owners further reinforces the functional convergence driven by cohabitation, potentially influencing host health via shared metabolic outputs.

Our findings provide critical mechanistic insights into the potential coevolutionary dynamics between human and pet (e.g., dog) oral cavity microbiomes, a process shaped by millennia of shared living environments, dietary overlap, and direct physical contact. The detected HGT events between canine and human oral cavity microbiota indicate a bidirectional functional integration of their microbiomes with profound implications for host disease susceptibility. For instance, transfer of xenobiotic degradation genes from dog microbiomes to human commensals has been shown to enhance the capacity of human oral cavity microbiota to metabolize environmental toxins and therapeutic agents, potentially altering drug bioavailability and efficacy [34]. Conversely, human-to-dog HGT of lipid metabolism-related genes may disrupt canine oral cavity microbial lipid homeostasis, leading to increased production of pro-inflammatory lipid mediators that could be transmitted to owners via close contact [32, 35]. Furthermore, the significant overlap in key metabolic pathways between human and pet microbiomes (including short-chain fatty acid synthesis, amino acid catabolism, etc) underscores the necessity of integrating pet microbiome data into human health research. For example, pet oral cavity microbiomes serve as a reservoir of functional genes involved in short-chain fatty acid production, which are critical for maintaining intestinal barrier integrity and regulating host immune responses. These mediators may be linked to impaired intestinal barrier function and systemic low-grade inflammation in humans, exacerbating metabolic dysfunction, etc.

Several limitations warrant consideration. First, the cross-sectional design prevents causal inferences about HGT directionality (e.g., dog-to-owner vs. owner-to-dog) and temporal dynamics. Second, functional annotations are dependent on database curation; updates to COG, eggNOG, or KEGG could alter pathway classifications and interpretations. Third, we did not validate HGT with direct evidence (e.g., PCR amplification of transferred genes, experimental functional assays), which is critical to confirm our hypothesis. Fourth, unmeasured confounders (such as dietary overlap between owners and pets or pet grooming practices) may have contributed to the observed functional similarities, potentially masking true HGT effects. Finally, sample size and demographic variability (e.g., age, breed, health status) were not fully described, which could limit the generalizability of our findings.

## Conclusion

In conclusion, our study provides compelling evidence for functional differentiation and convergence in the oral cavity microbiomes of dogs, owners, and volunteers, with implications for HGT and human-pet coevolution. Future studies addressing these limitations will be critical to advancing our understanding of the microbial interactions that shape shared human and pet health.

## Statements

## Author contributions

Chuantao Fang, Asadullah Abid: Writing – original draft, Formal analysis, Data curation. Shan Li, Yongxiang Li, Feng Liu, Lu Liu, and Zhuo Lan: Conceptualization, Resources. Guofeng Cheng: Writing – review & editing, Visualization, Supervision, Conceptualization, Funding acquisition, Resources.

## Acknowledgements

This study was supported by the Fundamental Research Funds for the Central Universities (Grant # 22120250298). We apologize for not being able to cite additional work owing to space limitations.

## Data availability statement

The sequencing data involved in the study will be available upon request.

## Ethics statement

The study was approved by the ethical committee of the School of Medicine of Tongji University (approved #: 2023tjdxsy031).

## Supporting information

**Fig. S1.**
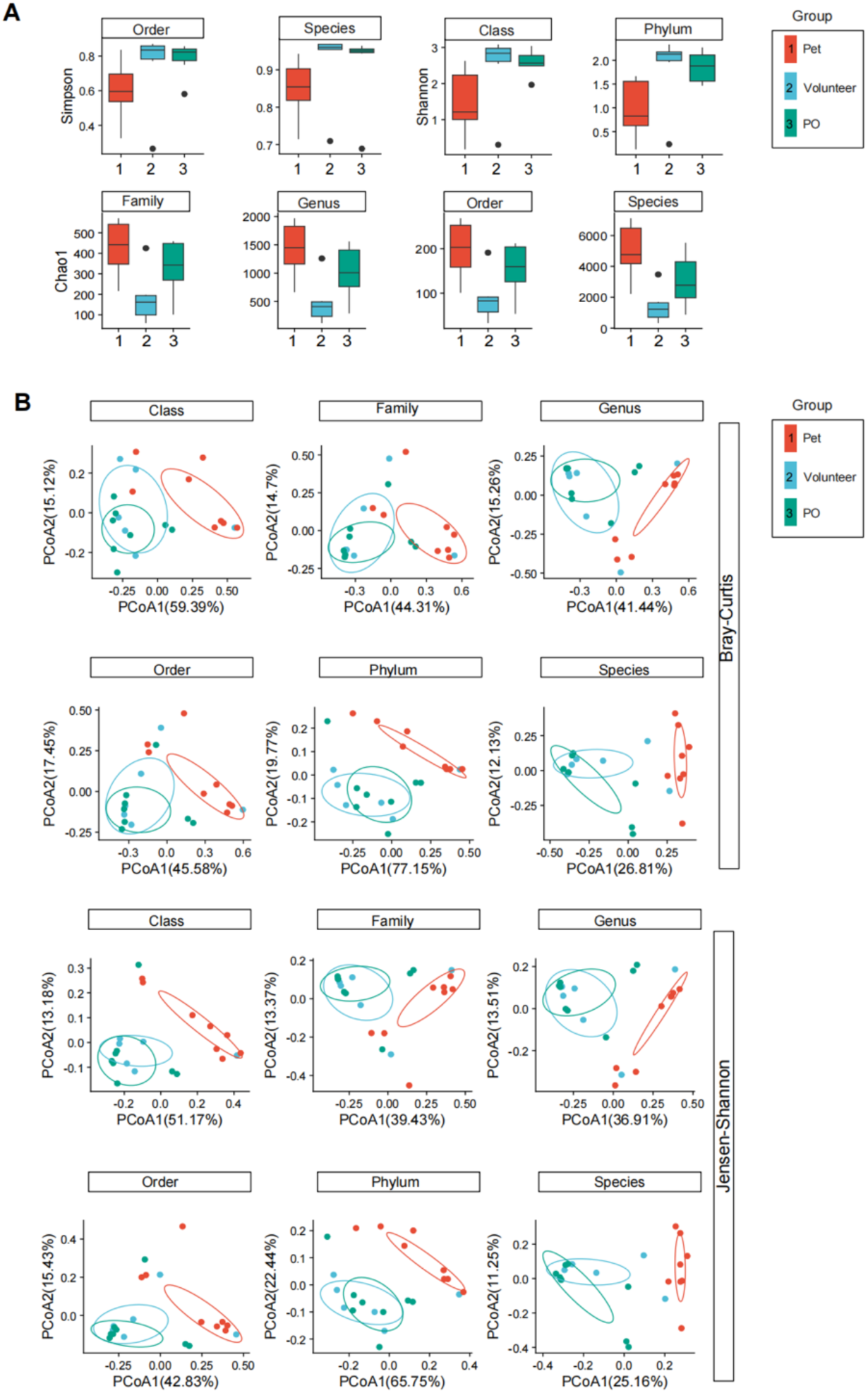
gene level analysis (related to Fig. 1) **A.** Chao1/Shannon/Simpson index representing the overall richness and evenness of microbial species across experimental groups. **B.** Principal Coordinates Analysis (PCoA) of metagenomic beta diversity. The PCoA was performed using Bray-Curtis and Jensen-Shannon distance, respectively.

**Fig. S2.**
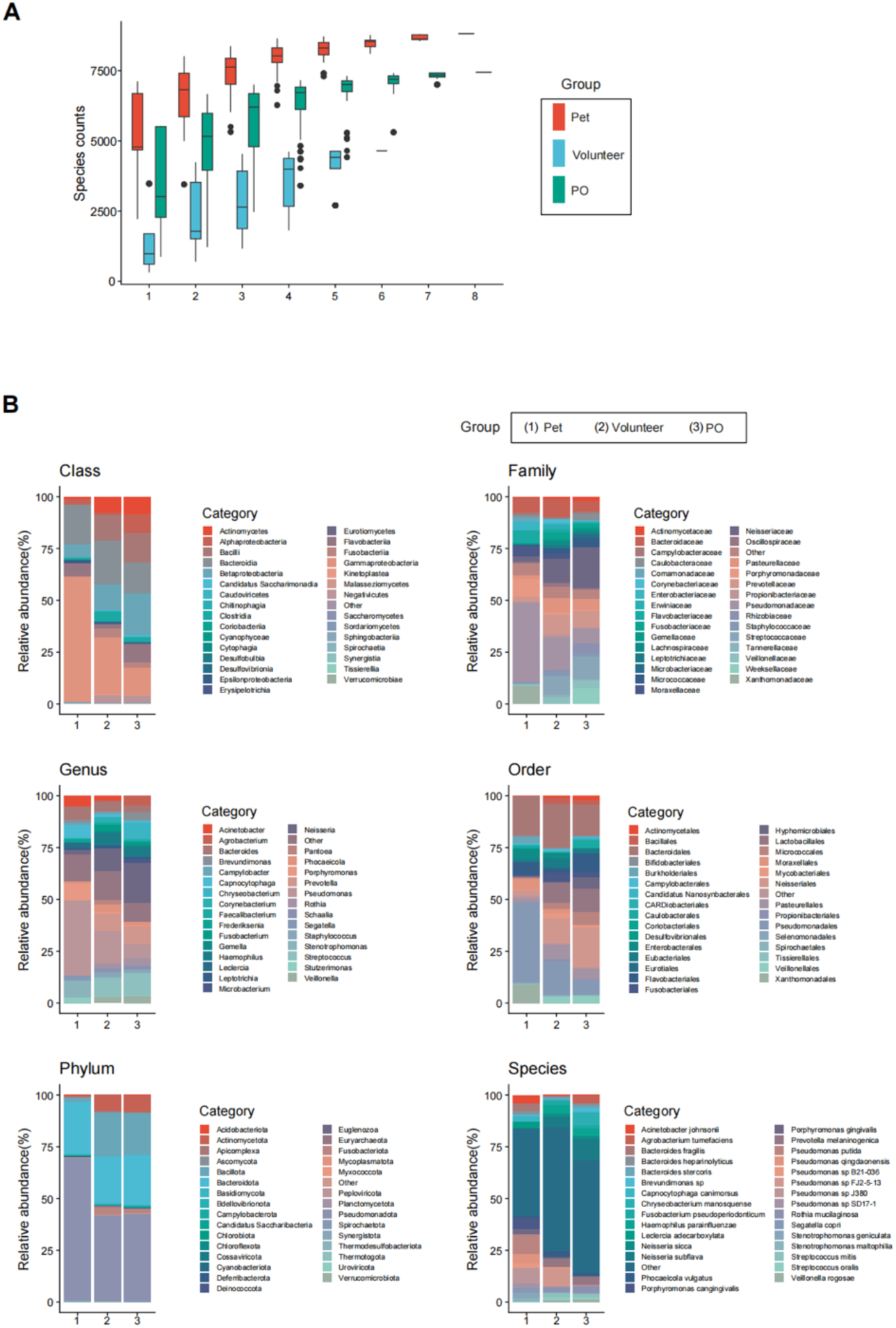
Taxonomy Analysis (related to Fig. 2). **A.** Boxplot showing the species counts of taxonomic components of Volunteer, PO (pet owner), and dog groups. PO, pet owner. **B.** Boxplot showing the relative abundance of taxonomic components of Volunteer, PO (pet owner), and dog groups. PO, pet owner. Taxonomic annotations were carried out at the Phylum, Class, Order, Family, Genus, and Species levels.

**Fig. S3.**
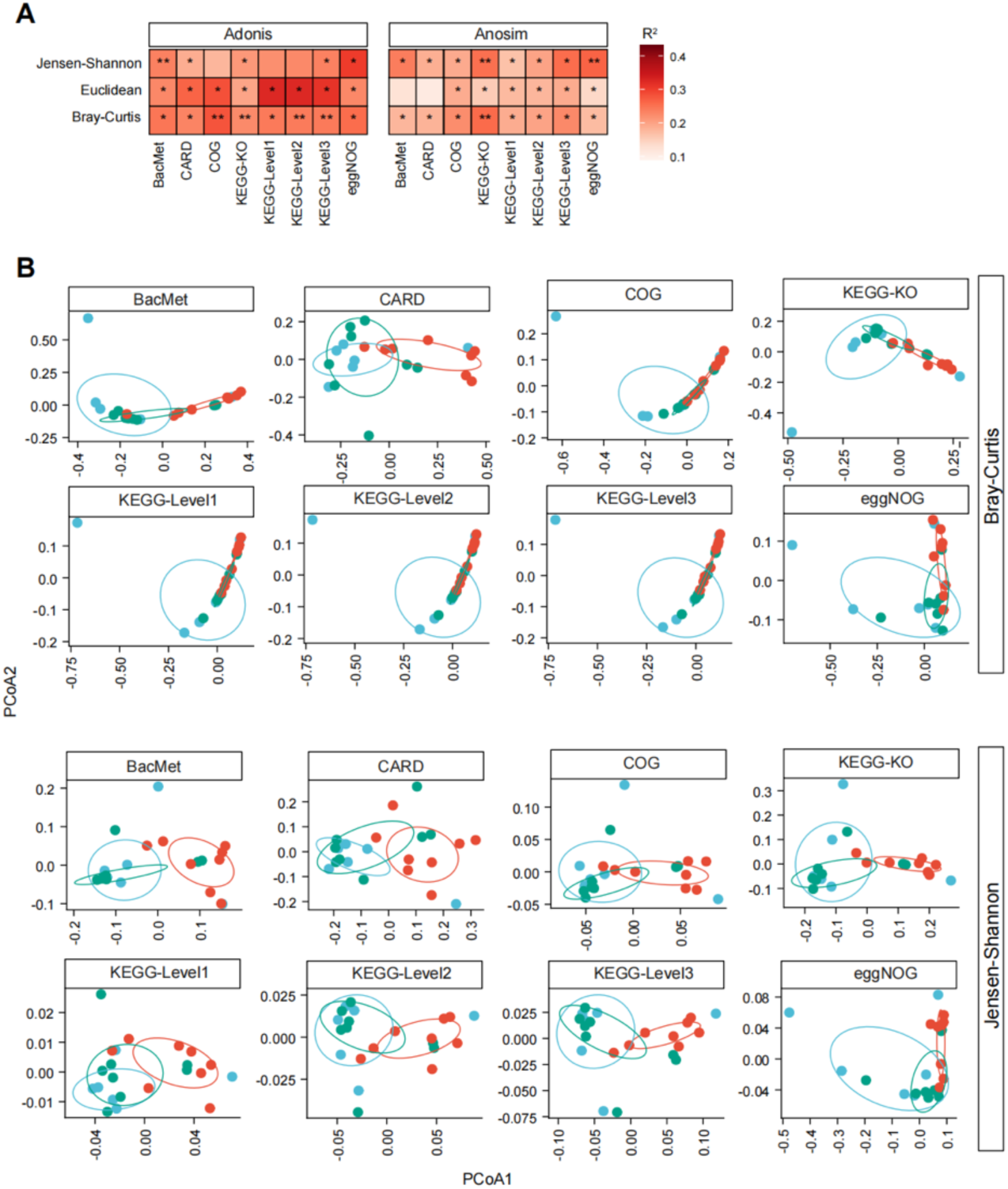
overview of functional annotation (related to Fig. 2). **A.** Heatmap showing the comparison results of beta-diversity of functional components among Volunteer, DO, and dog groups. The functional distance among samples in each group was reflected by the Bray-Curtis and Jensen-Shannon methods, respectively. Adonis and Anosim methods were used for statistical analysis. *p < 0.05, **p < 0.01, ***p < 0.001. **B.** Principal Coordinates Analysis (PCoA) of metagenomic beta diversity. The PCoA for the indicated functional components was performed using Bray-Curtis and Jensen-Shannon distance, respectively.

**Fig. S4.**
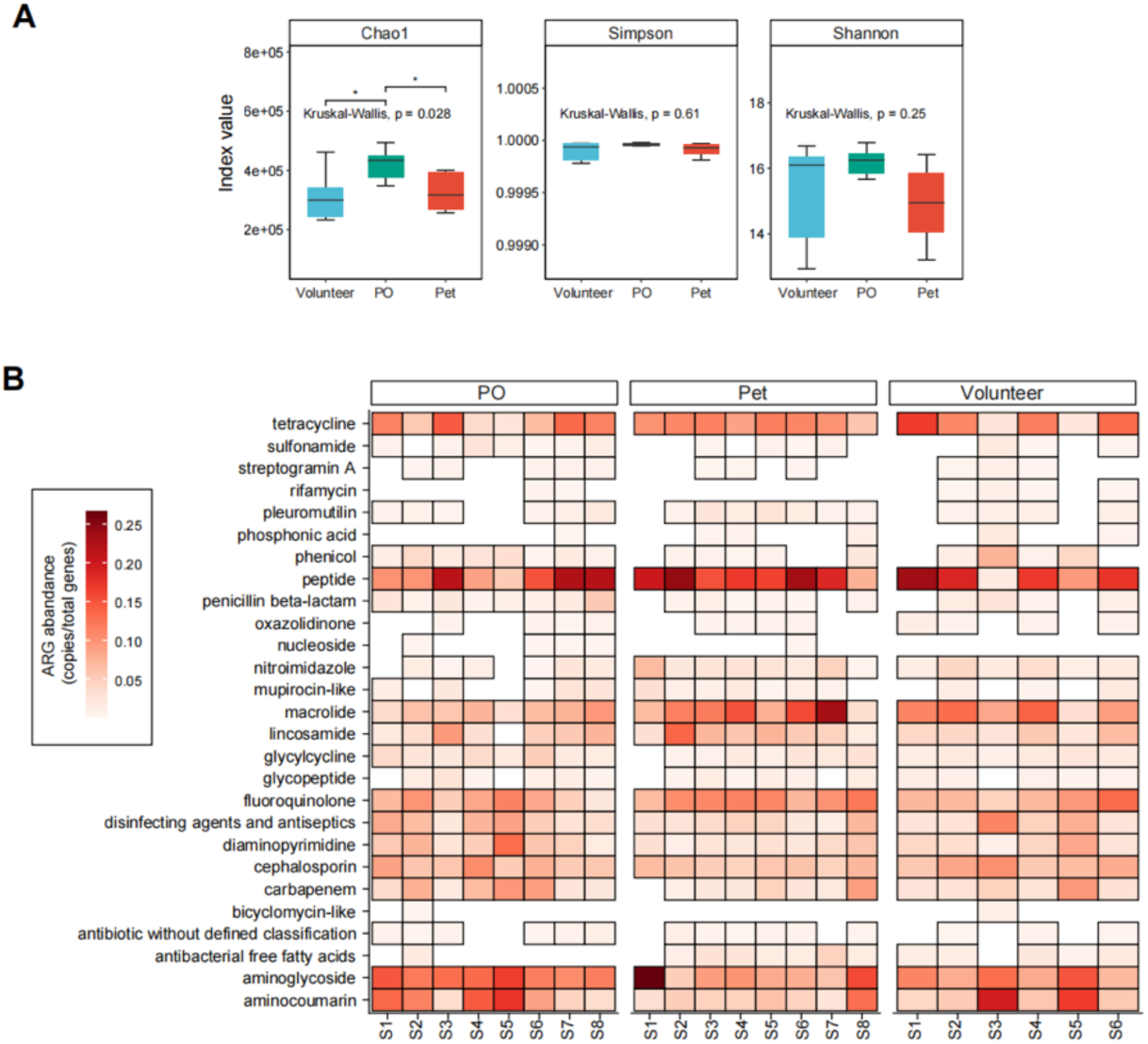
Comparison of resistome (ARGs) in the oral cavity of dogs and humans (related Fig. 3). **A.** Chao1/Shannon/Simpson index representing the overall richness and evenness of ARGs across experimental groups. ARGs, Antibiotic resistance genes. **B.** Heatmap showing the details of ARG components (per gene) of Volunteer, PO (pet owner), and dog groups. PO, pet owner.

**Fig. S5.**
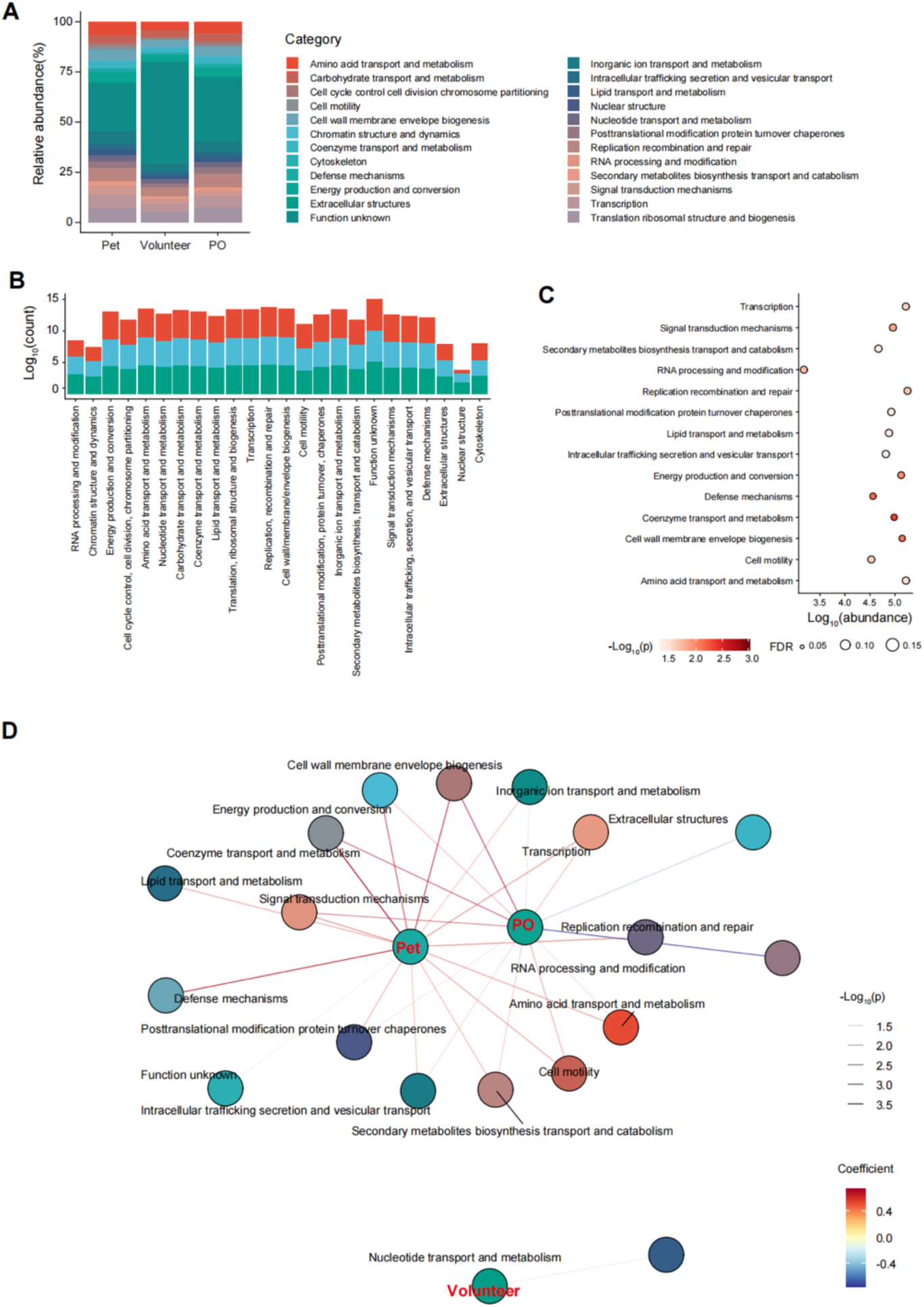
Cog /Nog annotation in the oral cavity of dogs and humans (related Fig. 5) **A.** Boxplot showing the relative abundance of eggNOG components of Volunteer, PO (pet owner), and dog groups. PO, pet owner; eggNOG, a public database of orthology relationships, gene evolutionary histories, and functional annotations. **B.** Barplot showing the total counts of eggNOG components of Volunteer, PO (pet owner), and dog groups. **C.** Dotplot showing the differentially expressed eggNOG components among Volunteer, PO (pet owner), and dog groups. **D.** The functional correlation analysis of eggNOG components.

**Fig. S6.**
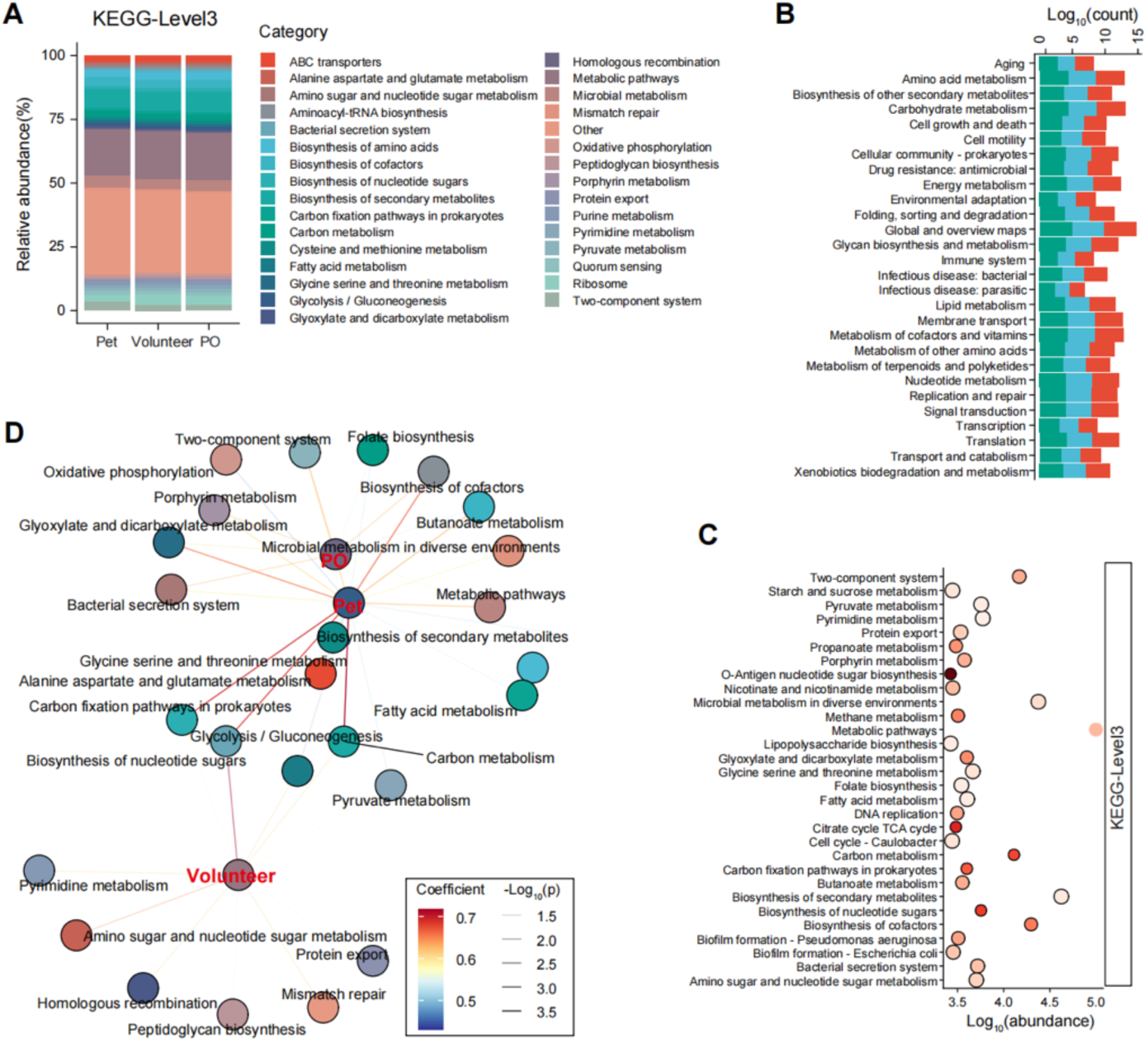
KEGG in the oral cavity of dogs and humans (related Fig. 6) **A.** Boxplot showing the relative abundance of KEGG (at level 3) components of Volunteer, PO (pet owner), and dog groups. PO, pet owner; KEGG, Kyoto Encyclopedia of Genes and Genomes. **B.** Barplot showing the total counts of eggNOG components of Volunteer, PO (pet owner), and dog groups. **C.** Dotplot showing the differentially expressed KEGG components (at level 3) among Volunteer, PO (pet owner), and dog groups. **D.** The functional correlation analysis at KEGG components (at level 3).

**Table S1** Quality control metrics for metagenome whole metagenome sequencing data using SOAPnuke and Bowtie2 aligner.

**Table S2** Detailed assembly statistics and quality metrics of whole metagenome sequencing data using MEGAHIT software.

**Table S3** The number of coding sequences annotated in the current study.

